# Peripheral inhibition of IL-6 signaling with tocilizumab improves stroke outcomes in aged mice but requires sex-specific dosing

**DOI:** 10.64898/2026.05.01.722347

**Authors:** Jacob Hudobenko, Eunyoung A. Lee, Gabi Delevati Colpo, Louise A. Atadja, Grant Goodman, Shuning Huang, Lucy Couture, Anjali Chauhan, Louise D. McCullough

## Abstract

Post-stroke inflammation contributes to poor outcomes in both clinical and experimental studies. Interleukin-6 (IL-6) is a key inflammatory mediator in ischemic stroke, and higher circulating IL-6 levels are associated with greater stroke severity and worse clinical outcomes. Targeting IL-6 signaling therefore represents a potential therapeutic strategy. We tested whether inhibition of IL-6 signaling with the IL-6 receptor (IL-6R) blocking antibody tocilizumab (TCZ) improves recovery after experimental stroke. Aged mice (18-20 months) underwent 60 minutes of middle cerebral artery occlusion. TCZ (20 mg/kg) was administered 5 hours after ischemia onset, and behavioral outcomes were assessed weekly for 5 weeks. Delayed TCZ treatment improved long-term functional recovery in aged male mice but not in aged females. To explore this difference, we measured circulating soluble IL-6R (sIL-6R) levels in mice and patients with ischemic stroke. Females exhibited significantly higher post-stroke sIL-6R levels. Increasing the TCZ dose to 100 mg/kg restored efficacy in aged female mice and improved long-term outcomes. These findings support a role for IL-6R pathway modulation in improving recovery after experimental stroke and suggest that therapeutic response may differ by sex and target availability, potentially related to differences in circulating sIL-6R after ischemic injury.

## Introduction

Stroke is a leading cause of long-term disability worldwide and disproportionately affects older adults. More than half of stroke survivors aged 65 years and older live with persistent functional impairment (1). Women carry a greater lifetime burden of stroke, with stroke ranking as the fifth leading cause of death in men and the third in women (2). Despite this, most experimental stroke studies continue to rely heavily on young male animals, and both preclinical and clinical studies frequently underreport outcomes by sex, despite NIH policies mandating their inclusion (3).

Intravenous thrombolysis remains the standard acute treatment for ischemic stroke but is limited by a narrow therapeutic window, strict eligibility criteria, and the risk of intracerebral hemorrhage (4). Tenecteplase offers practical advantages, including greater fibrin specificity and single-bolus administration, but remains similarly time dependent (5). Endovascular thrombectomy is highly effective for large vessel occlusion but requires specialized centers and remains inaccessible to many patients (6). These limitations highlight the need for therapies that target secondary injury pathways and improve recovery after stroke.

Aging and sex significantly influence stroke outcomes. Despite similar reductions in cerebral blood flow, aged mice have smaller infarcts but higher mortality and worse functional outcomes than young mice (7–9). In young animals, estrogen is associated with reduced injury and improved recovery in females compared to males (10). In contrast, aging is associated with a shift toward a more pro-inflammatory state, and clinical studies suggest that older individuals, particularly women, exhibit heightened inflammatory responses after stroke (11). These observations underscore the importance of incorporating both age and sex into experimental design and suggest that inflammatory pathways may contribute to differences in post-stroke recovery (11, 12).

Interleukin-6 (IL-6) is a pleiotropic cytokine produced by multiple cell types in response to injury (13). IL-6 signals through two distinct pathways. In classical signaling, IL-6 binds the membrane-bound IL-6 receptor (mIL-6R) and signals through gp130, restricting this pathway to cells that express mIL-6R (14–16). In contrast, trans-signaling occurs when IL-6 binds soluble IL-6 receptor (sIL-6R), enabling activation of gp130 on a broad range of cell types and amplifying inflammatory responses (14, 15, 17). While classical signaling is often associated with homeostatic or tissue-protective effects, trans-signaling has been linked to pathogenic inflammation and adverse outcomes after tissue injury. These distinct signaling pathways highlight sIL-6R as a key regulator of IL-6-mediated inflammatory response (15, 16).

The effects of IL-6 signaling in stroke are complex and context dependent and have been examined in numerous experimental studies (18). IL-6 contributes to both early and sustained inflammatory responses and may also exert neurotrophic effects (19). Importantly, its actions within the central nervous system may differ from those in the periphery. Locally produced IL-6 promotes angiogenesis after stroke (20) and intracerebroventricular administration of IL-6 is neuroprotective in permanent ischemia models in rats (21), supporting a potentially beneficial role for central IL-6 signaling. In contrast, circulating IL-6 levels increase with age and are associated with larger infarcts and worse outcomes after stroke (22, 23). Together, these observations suggest that selectively modulating peripheral IL-6 signaling while preserving potentially beneficial central IL-6 activity may represent a useful therapeutic strategy.

Tocilizumab (TCZ) is a humanized monoclonal antibody against IL-6R that is FDA-approved for the treatment of rheumatoid arthritis, giant cell arteritis, and cytokine release syndrome (24). TCZ blocks IL-6 signaling through both membrane-bound and soluble IL-6 receptors (25). In clinical studies, TCZ increased myocardial salvage and reduced neutrophil counts in patients with ST-segment elevation myocardial infarction (STEMI) (26, 27) and reduced systemic inflammation and myocardial injury in comatose patients resuscitated after out-of-hospital cardiac arrest (28, 29). More recently, a phase 2 clinical trial in ischemic stroke patients undergoing endovascular treatment reported that TCZ reduced infarct growth at 72 hours without increasing hemorrhagic transformation or other adverse events (30). Despite these findings, the role of IL-6 signaling in stroke recovery remains incompletely understood, particularly in the aging brain. Circulating IL-6 levels increase with age and are associated with worse outcomes after stroke, supporting a role for IL-6 in adverse post-stroke outcomes. We therefore hypothesized that peripheral inhibition of IL-6 signaling would improve post-stroke recovery. To test this, we evaluated the effects of delayed administration of tocilizumab in aged male and female mice following ischemic stroke.

## Results

### Delayed TCZ treatment (20 mg/kg) improves survival and functional recovery after stroke in aged male mice

While pretreatment with TCZ mitigates ischemia-reperfusion injury in rats, this approach has limited translational relevance for acute stroke therapy (31). We therefore evaluated whether delayed TCZ administration improves outcomes after stroke. TCZ (20 mg/kg, i.p., single dose) or an IgG control was administered five hours after ischemia onset in male and female mice (Fig. 1A). Pharmacokinetic studies indicate that TCZ has a prolonged half-life, ranging from 6 to 9 days in rats and monkeys and approximately 10 days in humans (32), supporting biweekly or monthly dosing in clinical use (32). Delayed TCZ administration significantly reduced hemispheric infarction in young male mice three days after MCAO, corresponding to the peak inflammatory period (Fig. 1B). We next confirmed that delayed TCZ treatment similarly reduced infarct volume in aged male mice at three days after MCAO (Fig. 1C).

**Figure 1.**
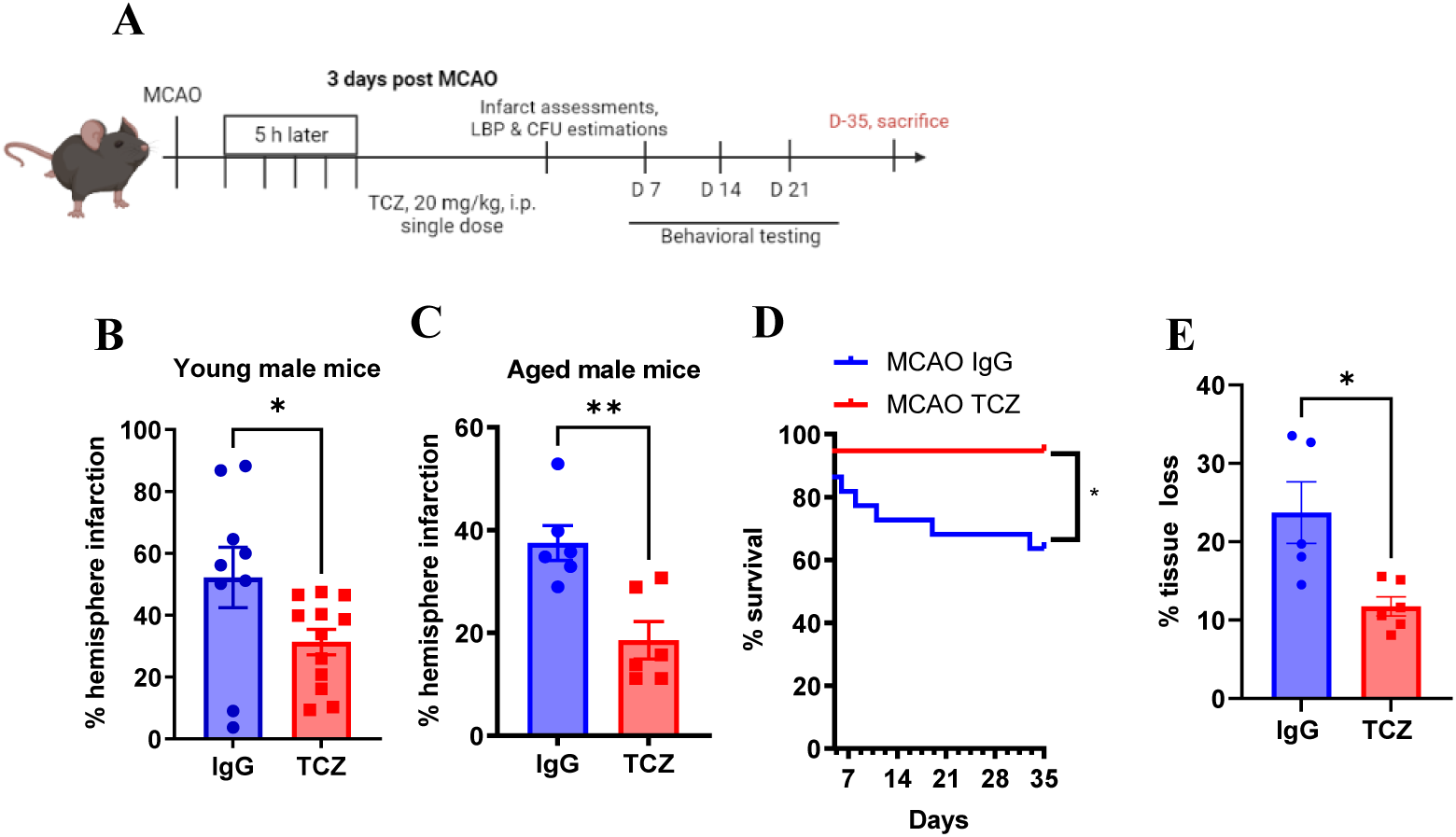
Delayed TCZ treatment (20 mg/kg) improves survival and reduces infarct size and atrophy in aged male mice. (A) Schematic of experimental design. (B) Percentage hemispheric infarction in young male mice at day 3 post-MCAO (n=9-12). (C) Percentage hemispheric infarction in aged male mice at day 3 post-MCAO (n=6). (D) Percentage survival at day 35 post-MCAO (n=19-22). (E) Percentage tissue loss at day 35 post-MCAO (n= 5-6). Data are presented as mean ± S.E.M. analyzed with the Mann-Whitney test. Two group comparisons were analyzed by unpaired t-test with Welch’s correction. *p<0.05, **p<0.01, ***p<0.001.

Because older patients have higher mortality rates and worse functional outcomes after stroke (34) we evaluated whether delayed TCZ improves survival and long-term recovery in aged mice. Aged male mice received TCZ (20 mg/kg, i.p.) five hours after MCAO and were followed for 35 days. Delayed TCZ significantly improved survival compared with IgG-treated controls (Fig. 1D). Functional outcomes were also improved. Neurological deficit scores (34) were significantly reduced from day 2 through day 6 (Fig. 2A). TCZ-treated aged males exhibited fewer right turns in the corner test on day 7, indicating improved sensorimotor symmetry (Fig. 2B). TCZ treatment also improved cognitive performance. In the Y-maze, TCZ-treated aged males exhibited significantly fewer direct revisits on day 7 (Fig. 2C). In the Barnes maze, TCZ-treated mice demonstrated higher escape rates compared with IgG controls (Fig. 2D). Delayed TCZ treatment also significantly reduced brain atrophy on day 35 after stroke (Fig. 1E).

**Figure 2.**
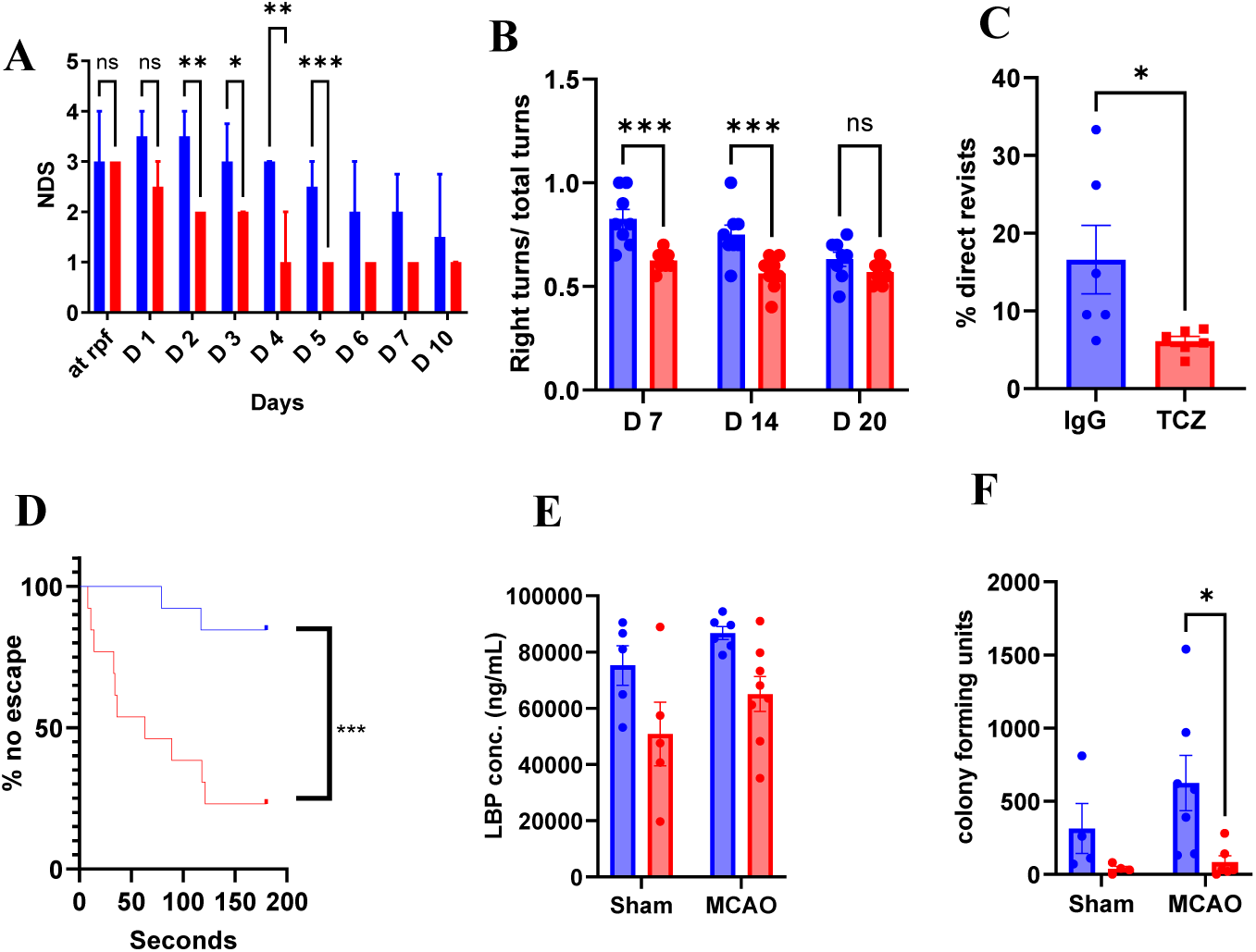
Delayed TCZ treatment (20 mg/kg) reduces bacterial burden and improves sensorimotor and cognitive outcomes in aged male mice after stroke. (A) Neurological deficit score (NDS; n=8). (B) Right turns in the corner test (n=8). (C) Percentage of direct revisits in the Y maze day 7 post-stroke (n=6). (D) Percentage of failed escape in the Barnes maze (n=13). (E) Plasma LBP concentration at day 3 post-MCAO (n=5=8). (F) Lung bacterial colony forming units at 35-days post-stroke (n=4-7). Data is presented as mean ± S.E.M. except NDS, which was presented as median (interquartile range) and was analyzed with the Mann-Whitney test. Two group comparisons were analyzed by unpaired t-test with Welch’s correction. Comparisons involving more than two groups were analyzed by two-way ANOVA with Sidak’s multiple comparisons test. *p<0.05, **p<0.01, ***p<0.001.

Because post-stroke infection contributes to morbidity and mortality (33) and TCZ modulates immune function, we assessed infection burden. Plasma lipopolysaccharide-binding protein (LBP) levels at 3-days post-stroke and lung bacterial colony-forming units (CFUs) were measured 35 days after MCAO. TCZ-treated mice had reduced lung CFUs compared with IgG-treated controls (Fig. 2F). Further we measured gut integrity 3-days post-stroke which did not reach significance but did demonstrate a trend in a reduced gut permeability post-stroke with TCZ treatment (Supp Fig 1). Together, these findings demonstrate that delayed TCZ treatment improves survival, reduces infection burden, and enhances long-term functional recovery after stroke in aged male mice.

### Delayed TCZ treatment (20 mg/kg) does not improve survival or functional recovery in aged female mice after stroke

Females remain underrepresented in both preclinical and clinical studies (3), contributing to variability in therapeutic responses and adverse drug effects (35). We therefore evaluated the effects of delayed TCZ treatment in aged female mice. In contrast to males, delayed TCZ administration (20 mg/kg, i.p.) did not improve survival compared with IgG-treated controls at 35 days after MCAO (Fig. 3A). Functional outcomes were also unchanged. Neurological deficit scores (Fig. 4A), corner test performance (Fig. 4B), Y-maze revisits (Fig. 4C), and Barnes maze escape rates (Fig. 4D) did not differ between TCZ- and IgG-treated mice, indicating no improvement in sensorimotor or cognitive function at this dose. Consistent with these findings, brain atrophy at day 35 after stroke was similar between groups (Fig. 3B). Measures of post-stroke infection were also unchanged, with no differences in plasma lipopolysaccharide-binding protein (LBP) levels or lung bacterial colony-forming units (CFUs) between TCZ- and IgG-treated females (Fig. 3C, D). Together, these findings indicate that TCZ at 20 mg/kg does not improve survival, histological injury, or functional recovery after stroke in aged female mice.

**Figure 3.**
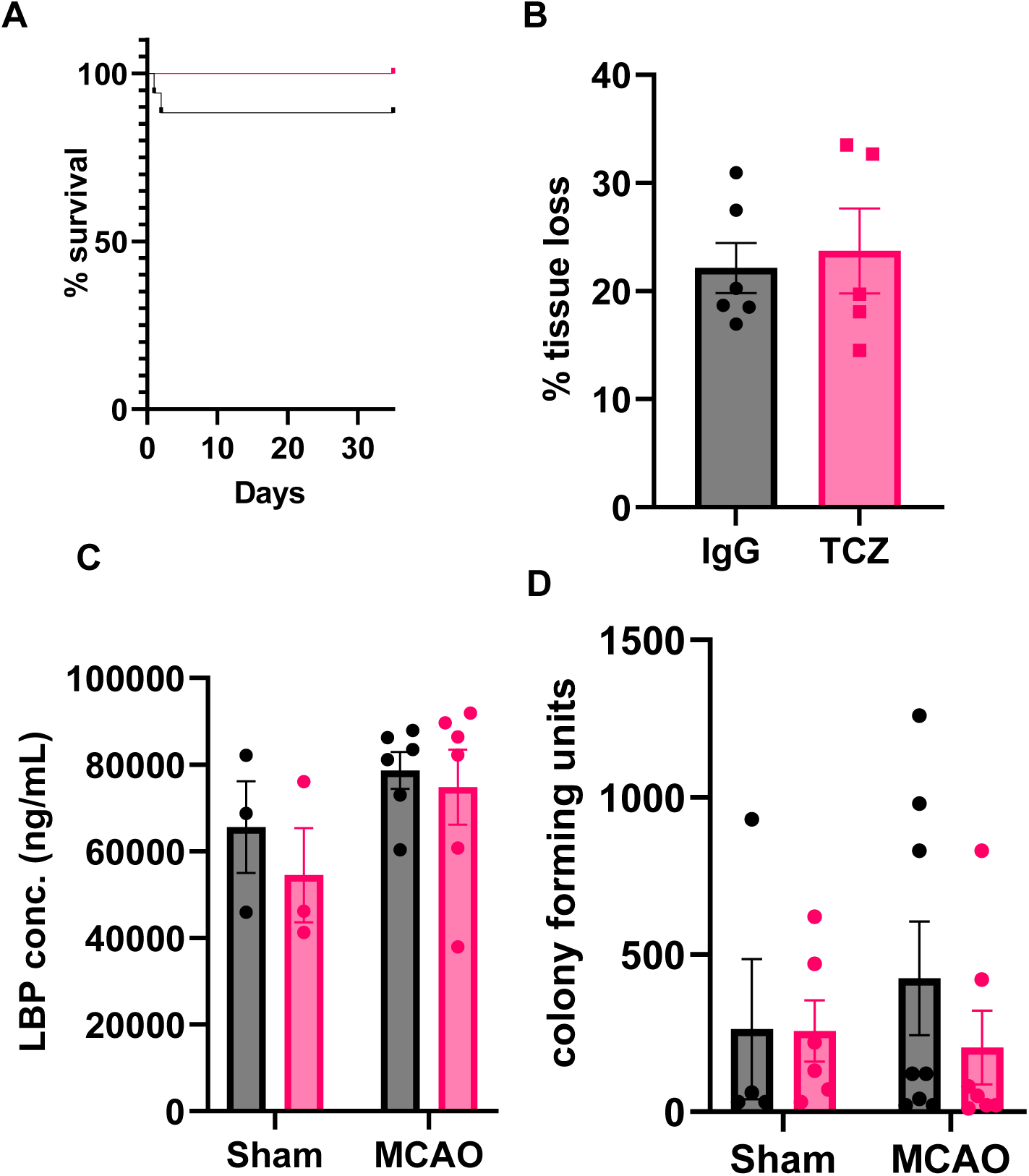
Delayed TCZ treatment (20 mg/kg) does not improve survival, brain atrophy, or bacterial burden in aged female mice after stroke. (A) Survival following MCAO (n=17). (B) Percentage tissue loss at day 35 post-MCAO (n=5-6). (C) Plasma LBP concentration 3-days post-stroke (n=3-6). (D) Lung bacterial colony-forming units 35-days post stroke (n=3-8). Data are presented as mean ± S.E.M. and analyzed with the Mann-Whitney test. Two group comparisons were analyzed by unpaired t-test with Welch’s correction. Differences among >2 groups were analyzed by Two-Way ANOVA with Sidak multiple comparisons.

**Figure 4.**
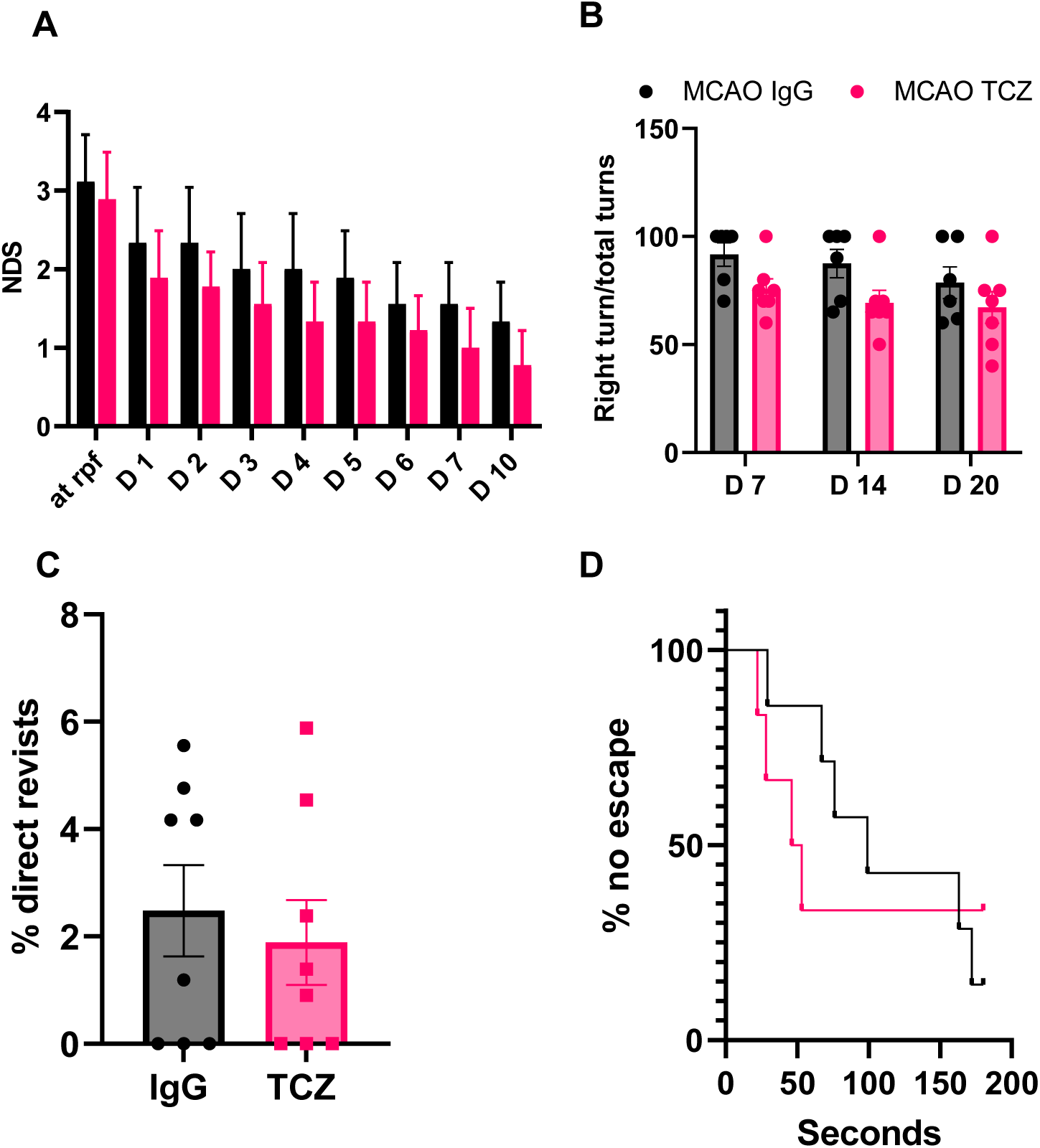
Delayed TCZ treatment (20 mg/kg) does not improve sensorimotor or cognitive outcomes in aged female mice after stroke. (A) Neurological deficit score (NDS; n=9). (B) Right turns in the corner test (n=6-7). (C) Percentage of direct revisits in the Y-maze day 7 post-stroke (n=8). (D) Percentage of failed escapes in the Barnes maze (n=6-7). Data are presented as mean ± SEM, except NDS, which is presented as median (interquartile range) and was analyzed using the Mann-Whitney test. Two-group comparisons were performed using unpaired t-tests with Welch’s correction. Comparisons involving more than two groups were analyzed by two-way ANOVA with Sidak’s multiple comparisons test.

### Aged females exhibit higher circulating sIL-6R levels after acute ischemic stroke

To investigate why TCZ was ineffective in aged females, we measured plasma soluble IL-6 receptor (sIL-6R) levels 24 hours after MCAO. Baseline sIL-6R levels did not differ between male and female sham mice (Fig. 5A). However, following MCAO, plasma sIL-6R levels were significantly higher in females compared with males (Fig. 5A).

**Figure 5.**
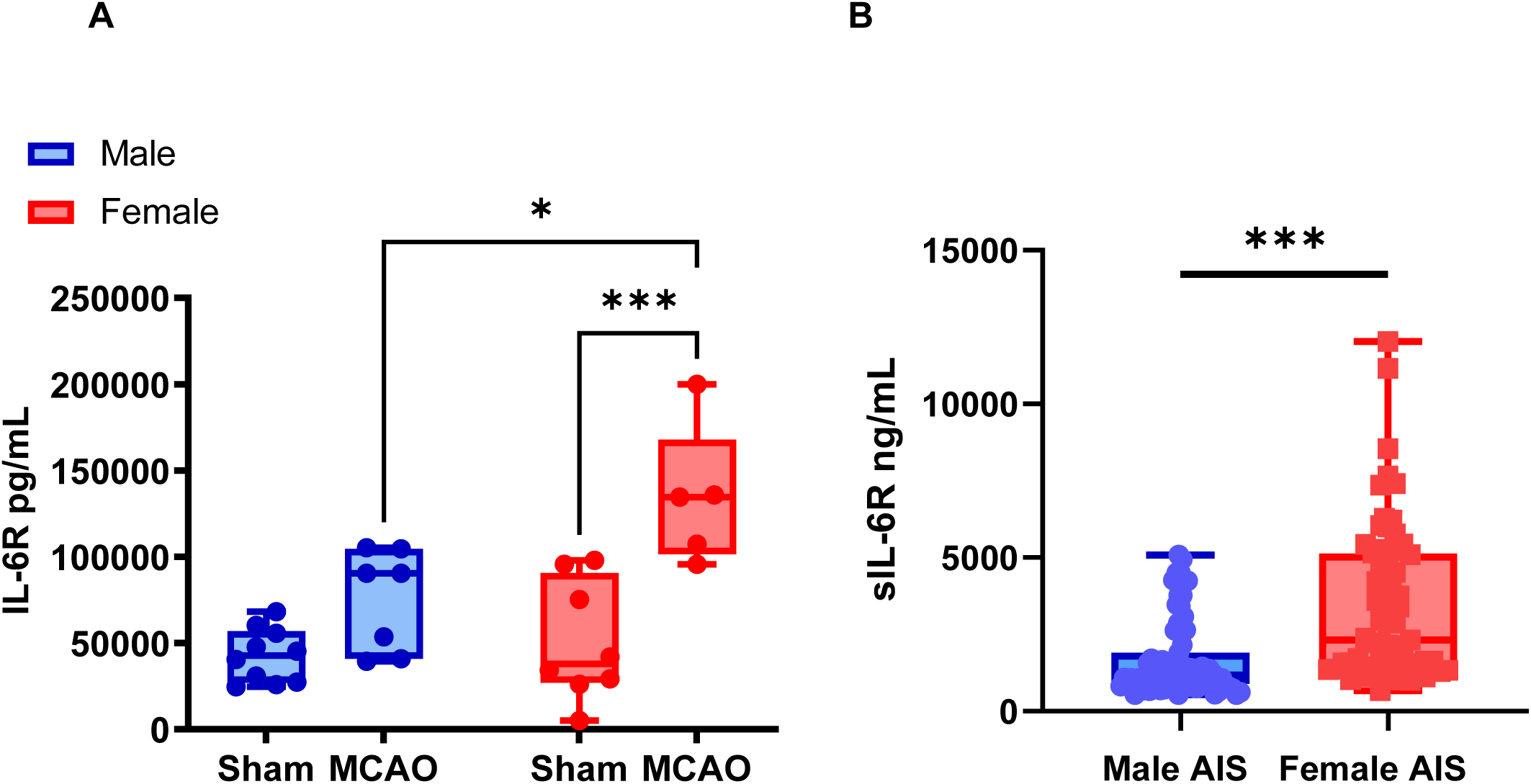
Aged female mice and women with acute ischemic stroke exhibit higher circulating sIL-6R levels. (A) Plasma sIL-6R levels in aged male and female mice 24 hours after sham or MCAO (n=5–10). (B) Plasma sIL-6R levels 24 hours after last known well in in patients with middle cerebral artery acute ischemic stroke (AIS) (n=67 females, n=71 males). Data in mice was analyzed using Two-Way ANOVA with Tukey’s multiple comparisons. Mouse data were analyzed using two-way ANOVA with Tukey’s multiple comparisons test. Human data were analyzed using univariable and multivariable linear regression on log-transformed values. *p<0.05, **p<0.01, ***p<0.001.

To determine whether this sex difference was also present in humans, we measured circulating IL-6, sIL-6R, and ADAM17, a metalloprotease involved in sIL-6R shedding (36), in patients with acute ischemic stroke. Women had significantly higher circulating sIL-6R levels after stroke compared with men (p<0.001; Fig. 5B). This association remained significant after adjustment for age and National Institutes of Health Stroke Scale (NIHSS) score in multivariable linear regression analysis (p<0.001; Supplementary Table 1B). In contrast, no significant sex differences were observed in circulating IL-6 or ADAM17 levels (Supplementary Table 1B). Together, these findings indicate that females exhibit higher systemic sIL-6R levels after stroke. This difference may influence the response to IL-6R blockade and suggests that higher dosing may be required to achieve effective target engagement in females.

### A higher TCZ dose (100 mg/kg) reduces infarction in female mice after stroke

We next tested whether increasing the TCZ dose could restore efficacy in females. A dose of 100 mg/kg was selected based on human dosing (8 mg/kg) scaled to mice using standard allometric conversion (×12.3). Aged male and female mice received TCZ (100 mg/kg, i.p.) five hours after MCAO, and infarct size was assessed three days later. In males, TCZ at 100 mg/kg reduced infarct size compared with IgG-treated controls (Fig. 6A) but did not provide additional benefit beyond the 20 mg/kg dose. In contrast, the higher dose significantly reduced infarct size in aged females, whereas the 20 mg/kg dose had no effect (Fig. 6B). Together, these findings indicate that higher TCZ dosing restores efficacy in aged female mice after stroke and is consistent with the higher circulating sIL-6R levels observed in females.

**Figure 6.**
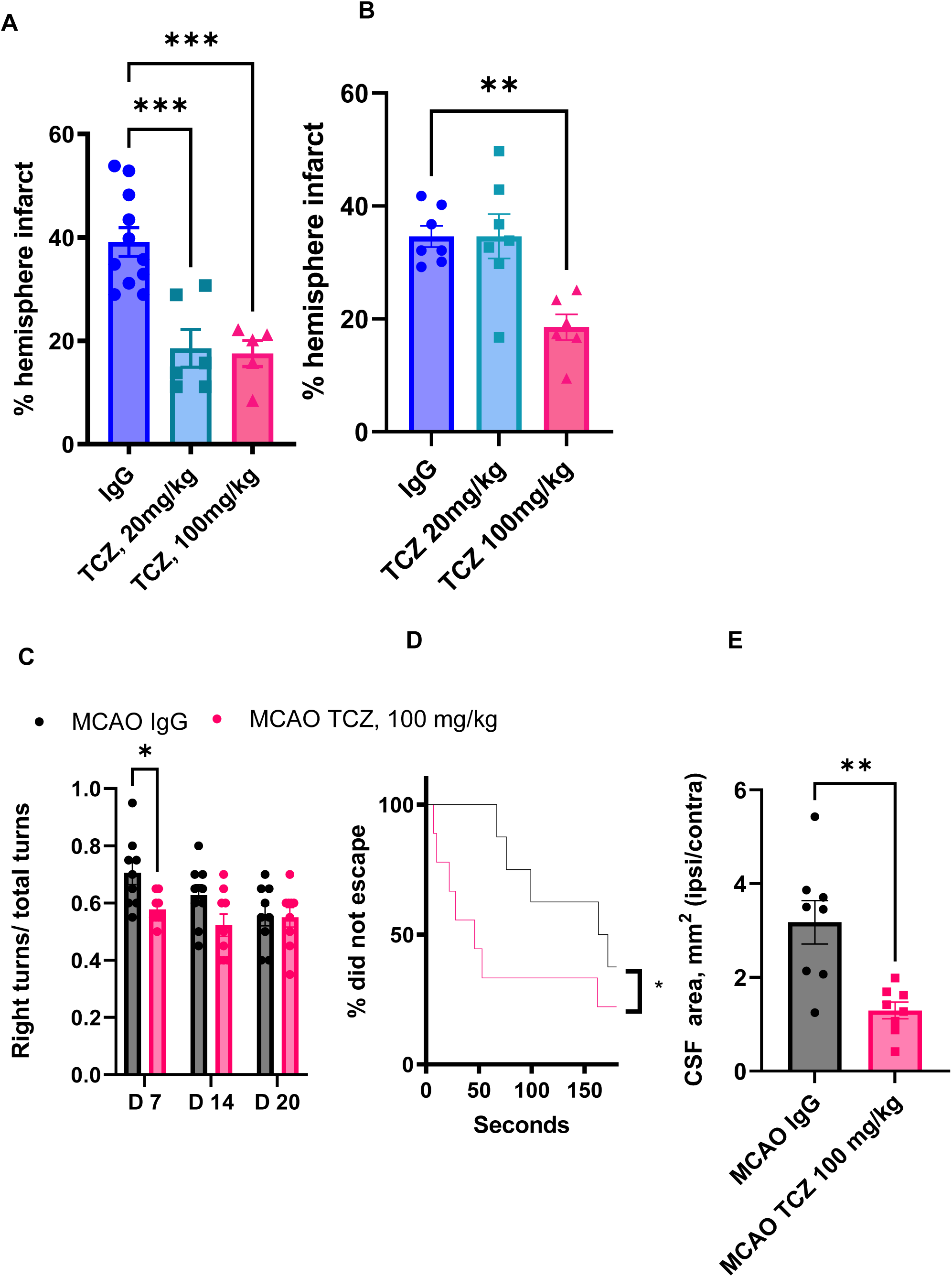
Higher-dose TCZ (100 mg/kg) reduces infarct size and atrophy and improves functional outcomes in aged female mice after stroke. (A) Percentage hemispheric infarction in males at day 3 post-MCAO (n=5-11). (B) Percentage hemispheric infarction in females at day 3 post-MCAO (n=6-7). C) Right turns in the corner test (n=9). (D) Percentage of successful escapes in the Barnes maze (n=8-9). (E) Female CSF area measurement of ipsilateral/contralateral sides as a measurement of tissue loss at day 35 post-MCAO measured using post-mortem brain MRI (n=8). Data are presented as mean ± SEM. Two-group comparisons were performed using unpaired t-tests with Welch’s correction. Hemispheric infarction was analyzed by one-way ANOVA with Dunnett’s multiple comparisons test. Corner test data were analyzed by two-way ANOVA with Sidak’s multiple comparisons test. *p<0.05, **p<0.01, ***p<0.001.

### Delayed high-dose TCZ improves cognitive outcomes and reduces cerebral atrophy in aged female mice after stroke

To determine whether higher-dose TCZ also improves long-term outcomes, aged female mice received a single 100 mg/kg dose five hours after MCAO. High-dose TCZ reduced right turns in the corner test (Fig. 6C) and improved Barnes maze escape performance (Fig. 6D), indicating improved sensorimotor and cognitive recovery. MRI analysis at day 35 showed reduced cerebrospinal fluid (CSF) signal area in TCZ-treated mice, consistent with reduced cerebral atrophy (Fig. 6E). Together, these findings indicate that higher-dose TCZ improves long-term functional outcomes and reduces cerebral atrophy in aged female mice after stroke.

### Delayed TCZ treatment has no effect in IL-6R knockout mice

To confirm that TCZ efficacy depends on IL-6R signaling, we administered TCZ (20 mg/kg) or IgG five hours after MCAO to young IL-6R⁻/⁻ mice (Fig. 7A, B). TCZ treatment did not alter infarct size in either male or female IL-6R⁻/⁻ mice at three days after stroke. These findings indicate that the neuroprotective effects of TCZ observed in wild-type mice depend on intact IL-6R signaling.

**Figure 7.**
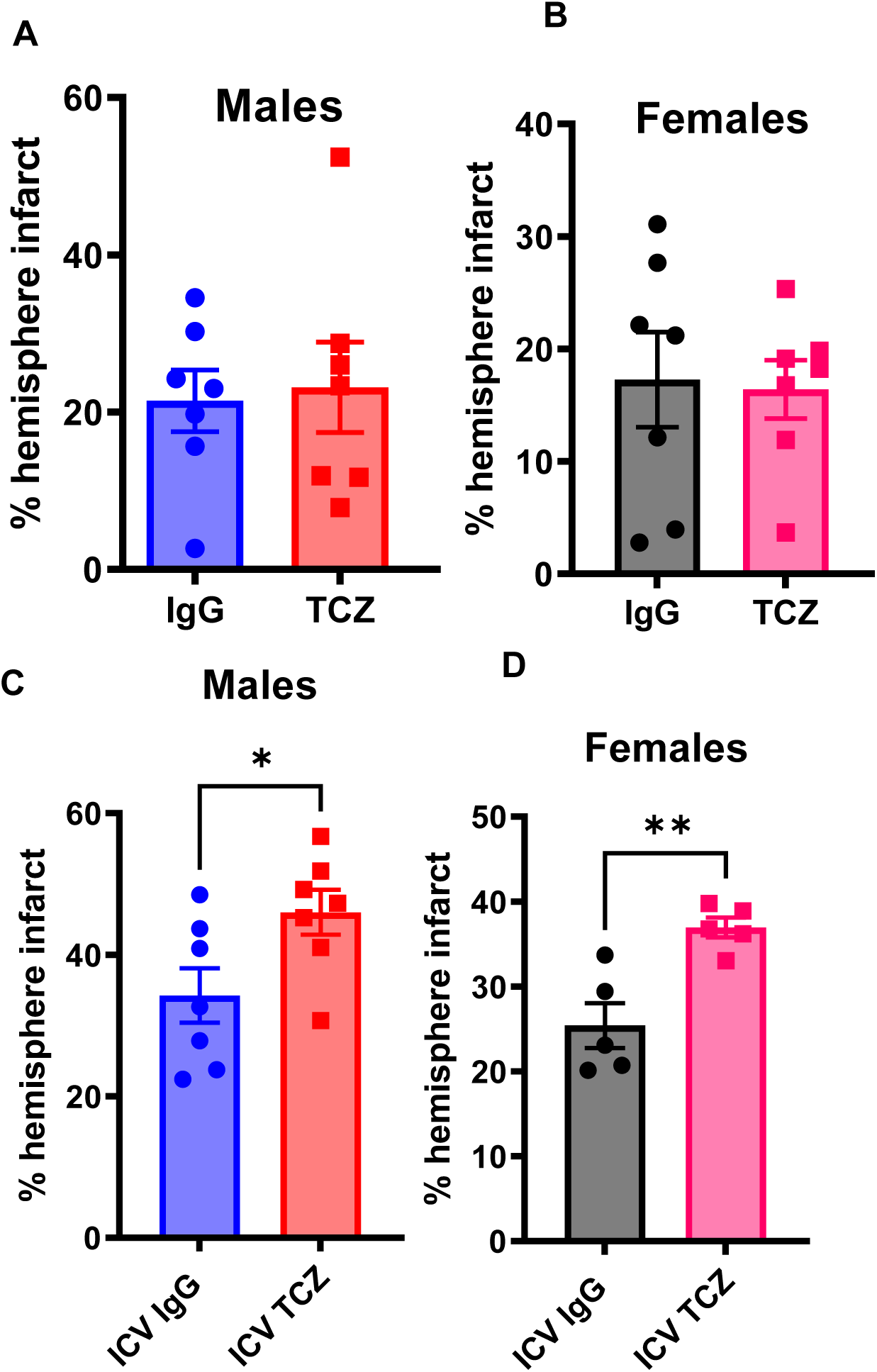
TCZ requires IL-6R signaling for efficacy, and central IL-6R inhibition worsens infarction after stroke. (A, B) Percentage hemispheric infarction in young IL-6R⁻/⁻ male (A) and female (B) mice at day 3 post-MCAO (n=7). (C) Percentage hemispheric infarction in young wild-type male mice following intracerebroventricular (ICV) TCZ administration at day 3 post-MCAO (n=7). (D) Percentage hemispheric infarction in young wild-type female mice following ICV TCZ administration at day 3 post-MCAO (n=5). Data are presented as mean ± SEM. Two-group comparisons were performed using unpaired t-tests with Welch’s correction. *p<0.05, **p<0.01, ***p<0.001.

### Intracerebroventricular administration of TCZ increases infarct size in wild-type mice

Prior studies suggest that inhibition of central IL-6 signaling may be detrimental after stroke, consistent with protective effects of IL-6 within the brain (37). To test this directly, we administered TCZ via intracerebroventricular (ICV) injection 12 hours before MCAO in young wild-type mice. ICV TCZ increased infarct size in both males (Fig. 7C) and females (Fig. 7D) compared with IgG-treated controls. These findings indicate that central IL-6 signaling is protective after stroke and support differential effects of IL-6 signaling in central and peripheral compartments.

### IL-6R shedding is linked to caspase-dependent signaling in female neutrophils

Previous studies have demonstrated sex differences in cell death pathways after experimental stroke, with female-derived cells preferentially activating caspase-dependent mechanisms, whereas male cells rely more heavily on caspase-independent pathways (38). Caspase inhibition is protective in females but not males in several stroke models, suggesting that apoptotic signaling plays a more prominent role in female ischemic injury (38). Because caspase activation can promote receptor cleavage and shedding, we hypothesized that sex differences in caspase signaling contribute to the elevated circulating sIL-6R levels observed in females after stroke.

To test this, we examined IL-6R expression and shedding in bone marrow-derived neutrophils from male and female mice. Neutrophils were selected because they are rapidly recruited after ischemic stroke and represent a major peripheral source of soluble IL-6R during inflammatory responses. Baseline IL-6R mRNA expression did not differ between male and female neutrophils (Fig. 8A). Following stimulation with 1 µM N-formyl-methionyl-leucyl-phenylalanine (fMLP), female neutrophils exhibited greater IL-6R shedding than male neutrophils. This effect was abolished by treatment with the pan-caspase inhibitor Q-VD-OPh (Fig. 8B), indicating that IL-6R shedding in female neutrophils is regulated by caspase-dependent mechanisms. Because receptor shedding is highly cell- and context-dependent, we next evaluated this relationship under ischemia-like stress. Splenocytes from male and female mice were subjected to oxygen-glucose deprivation (OGD) with or without caspase inhibition. In macrophages, caspase inhibition reduced IL-6R shedding and increased the proportion of IL-6R⁺ cells in both sexes (Fig. 8C), consistent with reduced receptor cleavage. In neutrophils, however, a sex-dependent pattern emerged: caspase inhibition increased IL-6R shedding in females but decreased shedding in males following OGD (Fig. 8D). These findings demonstrate that IL-6R shedding after ischemic stress is regulated by caspase-dependent pathways in a cell- and sex-specific manner and may contribute to the higher circulating sIL-6R levels observed in females after stroke

**Figure 8.**
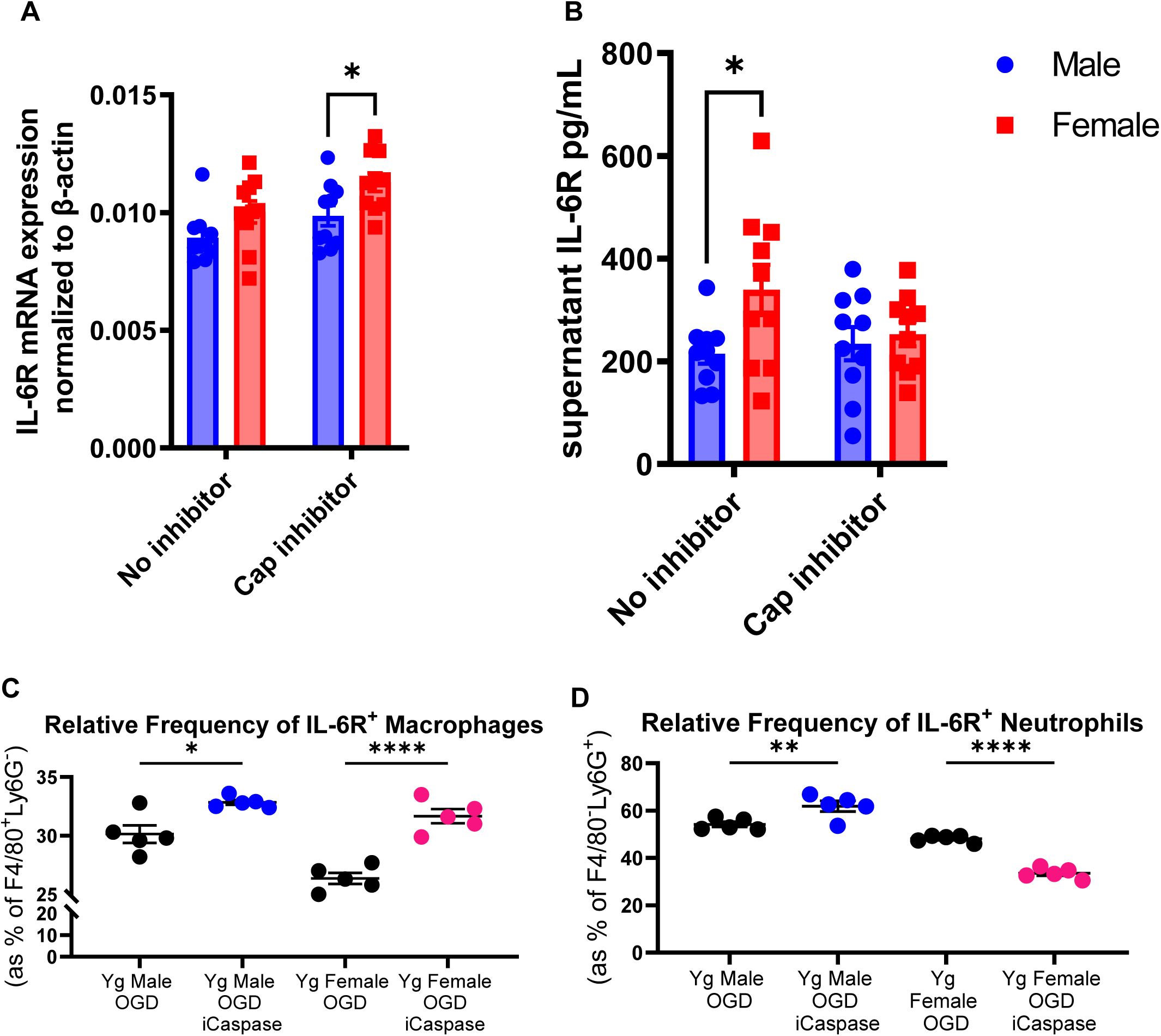
IL-6R shedding is cell type, sex, and caspase-dependent. (A) IL-6R mRNA expression in *ex-vivo* aged male and female neutrophils stimulated with 1 μM N-formyl-methionyl-leucyl-phenylalanine (fMLP), with or without caspase inhibitor (n=10). (B) Supernatant IL-6R levels in aged *ex-vivo* stimulated male and female neutrophils with or without caspase inhibitor (n=10). (C) Relative frequency of IL-6R⁺ macrophages (gated as F4/80⁺Ly6G⁻) following *ex-vivo* oxygen-glucose deprivation (OGD), with or without caspase inhibitor, in male and female splenocytes (n=4). (D) Relative frequency of IL-6R⁺ neutrophils (gated as F4/80⁻Ly6G⁺) following *ex-vivo* OGD, with or without caspase inhibitor, in male and female splenocytes (n=4). Data are presented as mean ± SEM. Panels A and B were analyzed by two-way ANOVA with Sidak’s multiple comparisons test. Panels C and D were analyzed by one-way ANOVA with Tukey’s multiple comparisons test. *p<0.05, **p<0.01, ****p<0.0001.

## Discussion

In this study, we demonstrate that delayed inhibition of IL-6 receptor signaling with tocilizumab (TCZ) improves survival, reduces infarct size, and enhances long-term functional recovery after ischemic stroke in aged mice. Notably, TCZ remained effective when administered five hours after ischemia, a clinically relevant therapeutic window. These protective effects were strongly sex- and dose-dependent. Delayed TCZ at 20 mg/kg improved survival and behavioral recovery in aged males but was ineffective in females, despite comparable injury. Increasing the dose to 100 mg/kg restored efficacy in females, improving cognitive outcomes and reducing cerebral atrophy. These findings identify IL-6R signaling as a modifiable pathway in post-stroke injury and demonstrate that therapeutic efficacy is determined by both biological sex and target burden. Consistent with prior studies demonstrating sex differences in pharmacologic responses, including differences in pharmacokinetics and pharmacodynamics that influence efficacy and toxicity (39, 40), our data indicate that IL-6R blockade is neuroprotective after stroke but requires sex-specific dosing.

The observed effects are consistent with modulation of IL-6R signaling. TCZ had no effect in IL-6R knockout mice, indicating that its protective actions require intact IL-6R. In contrast, intracerebroventricular administration of TCZ increased infarct size in both sexes, demonstrating that inhibition of central IL-6 signaling is detrimental. These findings define a compartment-specific role for IL-6 signaling after stroke. While elevated systemic IL-6 levels are associated with worse outcomes in patients, experimental studies indicate that IL-6 produced within the brain promotes angiogenesis and neuroprotection (20, 21). IL-6 signaling occurs through both classical signaling via membrane-bound IL-6R and trans-signaling mediated by soluble IL-6R (sIL-6R), which enables activation of gp130-expressing cells that do not express IL-6R (15, 16). Trans-signaling has been shown to amplify inflammatory responses and has been implicated in vascular inflammation and ischemic injury (14). In this context, the divergent effects of systemic versus intracerebral TCZ administration are consistent with selective modulation of peripheral IL-6 trans-signaling, while preservation of central IL-6 activity may be required for tissue repair. These findings suggest differential roles of peripheral and central IL-6 signaling after stroke, although the precise contribution of compartment-specific IL-6 activity will require further study.

The present findings extend our understanding of IL-6 signaling contributions to stroke pathology. IL-6 signals through both classical and trans-signaling pathways. In classical signaling, IL-6 binds to membrane-bound IL-6R, whereas in trans-signaling IL-6 interacts with soluble IL-6R (sIL-6R) to activate cells that do not express the membrane receptor (15,16). Trans-signaling amplifies inflammatory responses and has been strongly implicated in vascular inflammation, endothelial dysfunction, and ischemic injury (14). In particular, IL-6 trans-signaling promotes endothelial activation, cytokine production, and pro-thrombotic signaling through gp130-dependent pathways (14) and has been linked to hypoxia-associated vascular injury via STAT3 and HIF1α signaling (41). In our study, both aged female mice and women with acute ischemic stroke exhibited significantly higher circulating levels of sIL-6R post-stroke. Elevated sIL-6R increases the capacity for IL-6 trans-signaling, thereby amplifying systemic inflammatory responses following ischemic injury. This is consistent with clinical data demonstrating that IL-6 signaling correlates with infarct size, perfusion deficits, and adverse outcomes (18, 23), as well as established sex differences in inflammatory responses with aging (10,11). Shedding of IL-6R is mediated by ADAM proteases (36), providing a plausible mechanism for the increased circulating sIL-6R observed in females. Importantly, higher circulating receptor levels increase the amount of antibody required to effectively neutralize IL-6 signaling. This likely explains why a higher dose of TCZ was required to achieve benefit in females. Consistent with this, TCZ at 100 mg/kg, selected using standard allometric scaling (42), was effective in aged females, improving long-term cognitive outcomes and reducing cerebral atrophy. No additional benefit was observed in aged males at this higher dose, suggesting a plateau effect and supporting sex-specific differences in IL-6 signaling dynamics, receptor availability, and therapeutic responsiveness. While our findings support a role for elevated sIL-6R in enhancing IL-6 signaling in females, these studies do not directly establish causality between sIL-6R levels and therapeutic dose requirements. These findings position circulating sIL-6R as both a mechanistic mediator and a potential biomarker of target engagement in IL-6-directed therapies.

Sex differences in pharmacologic responses are well recognized and can arise from differences in drug metabolism, immune activation, and target availability (39, 40, 43). In parallel, sex differences in inflammatory signaling pathways are increasingly appreciated, and our data suggest that elevated sIL-6R in females reflects differences in IL-6R shedding. Prior work has demonstrated that cell death pathways diverge by sex after ischemia, with female cells preferentially engaging caspase-dependent apoptotic pathways, whereas male cells rely more heavily on caspase-independent mechanisms (38).

Consistent with this framework, we found that IL-6R shedding from neutrophils is regulated by caspase activity in a sex-dependent manner. Female neutrophils exhibited greater IL-6R shedding following stimulation, and this response was attenuated by caspase inhibition, supporting a role for caspase-dependent receptor cleavage. These findings provide a mechanistic link between sex-specific cell death signaling and regulation of IL-6R availability. Increased shedding of IL-6R would be expected to expand the circulating sIL-6R pool, thereby enhancing IL-6 trans-signaling and amplifying systemic inflammatory responses after stroke. This shift in ligand–receptor dynamics increases the effective target burden and may necessitate higher concentrations of TCZ to achieve sufficient receptor occupancy and pathway inhibition.

Our results provide important context for emerging clinical studies targeting IL-6 signaling in stroke. The recently published IRIS phase 2 clinical trial evaluated TCZ in patients with acute anterior circulation ischemic stroke undergoing endovascular treatment. TCZ-treated patients exhibited reduced infarct growth at 72 hours compared with placebo, without increased hemorrhagic transformation or serious adverse events (30), providing early clinical evidence that IL-6R inhibition can modify ischemic injury in humans. The IRIS trial did not report clear sex-specific differences in treatment response. This likely reflects limited power rather than true biological equivalence, given the modest sample size and imbalanced sex distribution across treatment groups. Women were underrepresented in the TCZ arm, and the overall cohort was not designed to detect sex-dependent differences in efficacy. Our findings suggest that such differences may be clinically relevant. Elevated circulating sIL-6R in females, as observed in both our preclinical and human cohorts, would be expected to influence target engagement and therapeutic response. These data support the incorporation of sex as a biological variable in future trials of IL-6R blockade and highlight the importance of integrating biomarker-based approaches, including sIL-6R and related inflammatory mediators, to define target engagement and optimize dosing strategies. These considerations may be particularly important in older populations, where inflammatory tone and IL-6 signaling are further amplified.

Our findings provide important context for prior results from the Stroke Preclinical Assessment Network (SPAN), in which TCZ advanced through initial screening but did not demonstrate consistent efficacy across models. In that platform, post-stroke mortality was high, approximately 42% in both treatment and control groups (44, 45), which can complicate interpretation of functional outcomes and reduce statistical power to detect treatment effects. Differences in experimental design, including variability in infarct size and mortality, may contribute to the differing conclusions regarding TCZ efficacy and highlight the importance of model selection in preclinical therapeutic evaluation. All experiments in the present study were randomized and performed with investigators blinded to treatment group; however, as with all preclinical studies, unrecognized sources of bias cannot be entirely excluded. At the same time, variability across preclinical platforms underscores the need for cautious interpretation, and further studies will be important to define the conditions under which IL-6R blockade is most effective.

An important limitation of this study is the species specificity of TCZ. Structural differences between human and murine IL-6R have led to concerns that TCZ may not fully inhibit murine IL-6 signaling (46), and preclinical development therefore relied on surrogate antibodies targeting murine IL-6R. Despite this, several lines of evidence indicate that TCZ achieved biologically meaningful modulation of IL-6R signaling *in vivo*. First, TCZ had no effect in IL-6R knockout mice, demonstrating that its activity requires the presence of IL-6R and is not mediated through off-target mechanisms. Second, we observed a clear dose–response relationship, with higher doses required in females exhibiting elevated circulating sIL-6R levels, consistent with concentration-dependent target engagement. Third, TCZ treatment produced a phenotype consistent with attenuation of IL-6 signaling reported in prior studies, including reduced infarct size, modulation of systemic inflammation, and improved functional recovery. These observations support the conclusion that, although TCZ likely has lower affinity for murine IL-6R compared with the human receptor, systemic administration at sufficient concentrations results in functionally relevant IL-6R blockade *in vivo* sufficient to modulate post-stroke inflammatory responses.

Post-stroke infections are a major clinical complication that intersects with inflammatory signaling pathways. Approximately one third of stroke patients develop infections, which are strongly associated with worse outcomes (33). Because TCZ modulates immune function, infection risk is an important consideration. In our study, TCZ treatment reduced pulmonary bacterial burden after stroke. This effect is most consistent with reduced stroke severity rather than a direct antimicrobial action, as the extent of brain injury is a primary determinant of post-stroke immunosuppression and infection risk (33). Further we measured gut integrity and plasma LBP levels 3-days post-stroke which did not reach significance but did demonstrate a trend in a reduced gut permeability and LBP levels post-stroke with TCZ treatment (Fig 2E and Supp Fig 1) which could help explain reduced CFU counts in TCZ treated mice. We are currently exploring the possible origins of these infections in subsequent studies. These findings suggest that modulation of IL-6 signaling may influence not only central injury but also downstream systemic complications following stroke. However, we cannot exclude direct immunomodulatory effects of IL-6R blockade on host defense, and the relationship between inflammation, stroke severity, and infection risk warrants further investigation.

In summary, delayed IL-6R inhibition with TCZ improves outcomes after experimental stroke in aged mice and highlights the context-dependent role of IL-6 signaling in ischemic brain injury. Some experiments, particularly in aged female cohorts, were performed with modest sample sizes and should be interpreted with appropriate caution. Our data support a model in which peripheral IL-6 signaling contributes to post-stroke inflammation and injury, whereas central IL-6 signaling may have protective effects. We further identify sex-dependent differences in IL-6R biology that influence therapeutic response, with elevated circulating sIL-6R in females increasing signaling capacity and necessitating greater receptor blockade to achieve efficacy. These findings have direct translational implications. The efficacy of IL-6-targeted therapies appears to depend not only on timing of administration but also on biological factors that determine target availability and pathway activation. The observation that females exhibit higher sIL-6R levels and require higher antibody dosing highlights the importance of incorporating biological sex and inflammatory target burden into therapeutic design. As clinical trials of IL-6R blockade in stroke continue to emerge, integration of mechanistic biomarkers such as sIL-6R may help guide dosing strategies and improve the likelihood of successful translation. These findings support a biomarker-guided approach to IL-6–targeted therapy, in which circulating sIL-6R levels may inform dosing and patient selection.

## Materials and Methods

Detailed descriptions of experimental procedures and materials are provided in the Supplementary Materials, including information on animal models, stroke induction, behavioral testing, gut permeability assessment, bacterial culture and colony-forming unit quantification, MRI acquisition and analysis (including CSF and ventricular measurements), ELISA assays and patient cohort characteristics, ex vivo neutrophil studies, oxygen-glucose deprivation experiments, flow cytometry, and statistical analyses.

## Acknowledgments

This work was funded by the R01NS128412 to AC and R35NS132265 to LDM. The authors would like to thank Ms. Yan Xu for maintaining the animal colony.

## Supplementary Materials

### Materials and Methods

#### Animals

Aged male and female C57BL/6J mice (18-20 months old; stock #000664, The Jackson Laboratory) were used for all primary experiments. A small preliminary cohort of young male C57BL/6J mice (3-4 months old) was also included. Young mice were purchased at approximately 2 months of age and acclimated for at least 4 weeks prior to experimentation. Aged mice were purchased at 6–9 months of age and maintained in the animal facility until reaching 18–20 months of age.

All animals were housed under specific pathogen-free conditions at McGovern Medical School with ad libitum access to food and water. All experimental procedures were conducted in accordance with NIH guidelines for the care and use of laboratory animals and were approved by the Institutional Animal Care and Use Committee at The University of Texas Health Science Center at Houston. Animals were randomly assigned to surgical and treatment groups.

#### IL-6R Knockout Mice

Interleukin-6 receptor (IL-6R) knockout mice were generated by crossing IL-6Rα floxed mice (stock #012944, The Jackson Laboratory) with CMV-Cre transgenic mice (stock #006054, The Jackson Laboratory) to achieve germline deletion of IL-6R.

Genotyping was performed by PCR using primers provided by The Jackson Laboratory to detect wild-type, floxed, and recombined IL-6Rα alleles. The following primers were used: wild-type IL-6Rα allele (5′-CTGGGACCCGAGTTACTACTT-3′ and 5′-CAGCAACACCGTGAACTCCTTT-3′), floxed IL-6Rα allele (5′-GCCTGGGTGGAGAGGCTTTTT-3′ and 5′-CCCAGTGAGCTCCACCATCAAA-3′), and recombined IL-6Rα allele (5′-CTGGGACAGGGAAGGGCTTTT-3′ and 5′-CCCAGTGAGCTCCACCATCAAA-3′).

Experimental knockout mice were identified by the absence of both wild-type and floxed IL-6Rα alleles and the presence of the recombined allele. Because recombination occurs in the germline, the CMV-Cre transgene was not maintained in experimental animals to avoid potential Cre-related effects.

#### Transient Stroke Model

Mice were randomly assigned to stroke or sham surgery and further subdivided into treatment groups, resulting in four groups per cohort. Transient cerebral ischemia was induced by 60 minutes of reversible middle cerebral artery occlusion (MCAO) as previously described (7). A 6-0 silicone-coated monofilament (Doccol Corporation, Sharon, MA) was introduced into the right internal carotid artery to occlude the origin of the middle cerebral artery. Following the occlusion period, mice were placed in a temperature-controlled recovery cage for 1 hour prior to reperfusion, at which time the monofilament was withdrawn to restore cerebral blood flow. Rectal temperature was continuously monitored throughout MCAO and reperfusion and maintained at 37°C. Sham-operated mice underwent identical procedures except that the monofilament was not advanced to occlude the middle cerebral artery.

Five hours after ischemia onset, mice received a single intraperitoneal injection of tocilizumab (20 mg/kg or 100 mg/kg for the high-dose cohort) or control human IgG (R&D Systems, Minneapolis, MN). Postoperatively, mice received daily subcutaneous injections of sterile saline and wet mash for 7 days following reperfusion. Animals were euthanized at either 3 or 35 days after MCAO

#### Drug Dosing

For the 35-day cohorts of aged male and female mice and the 3-day cohort of young male mice, animals received a single intraperitoneal injection of tocilizumab or control human IgG 5 hours after ischemia onset at doses of 20 mg/kg or 100 mg/kg.

#### Behavioral Testing

Behavioral assessments were performed to evaluate neurological, sensorimotor, cognitive, and exploratory outcomes following stroke. Neurological deficit scores were assessed using the Bederson 5-point standardized scoring (34), daily during the first week after surgery (days 1-7) and subsequently on days 10, 14, 20, 28, and 35. Sensorimotor asymmetry was evaluated using the corner test on days 7, 14, and 20 after surgery (48). Working memory and exploratory behavior were assessed using the Y-maze test on days 7 and 21 (49). These time points were selected to minimize repeated testing while enabling longitudinal assessment of recovery.

Spatial learning and memory were assessed using the Barnes maze. Training was conducted on days 27–29 after surgery, and probe testing was performed on day 31. The Barnes maze consisted of an elevated circular platform with 20 equally spaced holes, one of which led to an escape box located beneath the platform. Visual cues were positioned around the testing room to facilitate spatial navigation. At the start of each trial, mice were placed in a central start chamber for 1 minute. After removal of the chamber, mice were allowed to explore the maze for up to 3 minutes to locate the escape hole. If a mouse failed to locate the escape hole within the allotted time, it was guided to the correct location and allowed to remain there for 1 minute. Mice underwent three trials per day during training

Behavioral testing was performed prior to surgery to establish baseline performance and identify animals with preexisting bias. All behavioral testing and analysis were conducted by investigators blinded to treatment group. Behavioral videos were analyzed using EthoVision XT software (Noldus Information Technology, Leesburg, VA).

#### Cresyl Violet and TTC Staining

Cresyl violet staining was used to quantify infarct volume at 3 days and brain atrophy at 35 days after stroke. Mice were perfused with 1× PBS followed by 4% paraformaldehyde. Brains were post-fixed and cryoprotected in 30% sucrose at 4°C overnight. Coronal sections (30 μm) were cut using a microtome. Eight equally spaced sections per brain, separated by 360 μm, were mounted onto slides and stained with cresyl violet. Images were acquired and analyzed by an investigator blinded to treatment group using SigmaScan Pro software to quantify infarct area and tissue atrophy.

For 2,3,5-triphenyltetrazolium chloride (TTC) staining, animals were euthanized and brains were rapidly removed and briefly chilled at −80°C for 4 minutes to facilitate sectioning. Brains were cut into five 2-mm coronal sections spanning from the olfactory bulb to the cerebellum. Sections were incubated in 1.5% TTC (Sigma, St. Louis, MO) and subsequently fixed in 4% formalin. Digital images were acquired, and infarct area was quantified using SigmaScan Pro software.

#### Gut Permeability Assessment

Intestinal permeability was assessed 3 days after surgery using fluorescein isothiocyanate–dextran (FITC–dextran, 4 kDa). FITC–dextran was prepared at 50 mg/mL in sterile water. On day 3 after surgery, mice were fasted for 5 hours prior to testing and then administered FITC–dextran by oral gavage at a dose of 6 mg per 10 g body weight. One hour after gavage, mice were euthanized and blood was collected by cardiac puncture using heparinized syringes. Blood samples were centrifuged at 14,000 rpm for 14 minutes to isolate plasma. Plasma FITC–dextran concentrations were measured using an EnSpire plate reader (PerkinElmer) with excitation at 470 nm and emission at 520 nm. A standard curve was generated using serial dilutions of FITC–dextran, along with blank controls.

#### Lung Bacterial Colony-Forming Unit (CFU) Measurement

Pulmonary bacterial burden was assessed by quantifying colony-forming units (CFUs) in lung homogenates at 35 days after surgery. Following euthanasia, lungs were collected, weighed, and homogenized in sterile 1% saponin in 1× PBS at a concentration of 50 mg tissue per mL. An aliquot of 50 μL of homogenate (corresponding to approximately 2.5 mg of lung tissue) was plated onto 5% sheep blood agar plates (Carolina Biological Supply, Burlington, NC) and incubated at 37°C for 16 hours. After incubation, bacterial colonies were counted. CFU values were normalized to tissue weight and expressed as CFUs per mL of homogenate.

#### MRI Acquisition and Analysis

Magnetic resonance imaging (MRI) was performed using a Bruker 7T Avance system equipped with a 20-cm bore, microgradients, and ParaVision 5.1 software. Brain images were acquired using a rapid acquisition with relaxation enhancement (RARE) sequence with the following parameters: repetition time (TR) 2500 ms, echo time (TE) 36 ms, and number of averages (NA) 2. The field of view was 3 cm with a matrix size of 256 × 256 and a slice thickness of 500 μm. MRI analyses were performed in mice treated with the 100 mg/kg dose of tocilizumab.

#### Assessment of Cerebrospinal Fluid and Ventricular Volume

Following transcardial perfusion with PBS followed by 4% paraformaldehyde (PFA), brains were harvested with the skull intact. Skin, muscle, ears, nasal tip, and lower jaw were removed to expose the skull. The intact head was fixed in 4% PFA at 4°C and subsequently transferred to 40 mL of 0.01% sodium azide in PBS with gentle agitation for 7 days at 4°C. Samples were then incubated in a contrast solution containing 5 mM gadopentetate dimeglumine (Bayer HealthCare Pharmaceuticals Inc., Wayne, NJ) and 0.01% sodium azide in PBS for 21 days at 4°C to enhance MRI contrast.

MRI datasets were analyzed using OsiriX MD software. DICOM images were imported, and whole-brain segmentation was performed. Regions corresponding to cerebrospinal fluid (CSF) and ventricular spaces were identified on each slice. Ventricular and CSF areas ipsilateral and contralateral to the injury were manually delineated and quantified to assess brain atrophy and ventricular enlargement. All MRI segmentation and analyses were performed by investigators blinded to treatment group. Assessment of cerebrospinal fluid and ventricular volume.

#### ELISA

Plasma levels of IL-6, soluble IL-6 receptor (sIL-6R), and ADAM17 in human samples were measured using commercially available ELISA kits (R&D Systems, D6050B; Abcam, ab46029; and Invitrogen, EHADAM17, respectively) according to the manufacturers’ instructions.

For mouse plasma and neutrophil culture supernatants, IL-6 and sIL-6R levels were measured using ELISA kits from R&D Systems (DY008 and DY1830) according to the manufacturers’ protocols. Samples and standards were processed as specified by each kit, and absorbance was measured using a microplate reader.

#### *Ex-vivo* Neutrophil Experiments

Bone marrow neutrophils were isolated from mouse femurs following perfusion. Femurs were dissected, cleared of surrounding tissue, briefly sterilized in 70% ethanol, and rinsed in 1× PBS. Bone marrow cells were collected by flushing the femurs with 5 mL Hanks’ Balanced Salt Solution (HBSS) without calcium or magnesium, supplemented with 2% heat-inactivated fetal bovine serum (FBS).

The cell suspension was filtered through a 40 μm cell strainer into a 50 mL conical tube and washed twice with HBSS containing 2% FBS. Cells were resuspended and layered onto a discontinuous Histopaque gradient (Histopaque 1.077 and 1.119; Sigma). Samples were centrifuged at 500 × g for 30 minutes at room temperature with no brake. Mononuclear cells at the HBSS–Histopaque 1.077 interface were discarded, and neutrophils were collected from the interface between Histopaque 1.077 and 1.119. Cells were washed twice with HBSS containing 2% FBS and further purified using a Mouse Neutrophil Isolation Kit (STEMCELL Technologies, catalog #19762) according to the manufacturer’s instructions. Neutrophil purity was routinely >90%, as assessed by morphological criteria and differential cell counts.

Isolated neutrophils were plated in 6-well plates and treated with the pan-caspase inhibitor Q-VD-OPh (MP Biomedicals) or vehicle control. Cells were stimulated with 1 μM N-formyl-methionyl-leucyl-phenylalanine (fMLP; Sigma-Aldrich) for 1 hour. Supernatants and cell lysates were collected for ELISA and qPCR analyses.

#### Oxygen Glucose Deprivation (OGD) and Flow Cytometry

To model ischemia-reperfusion conditions *in vitro*, splenocytes isolated from male and female mice were subjected to oxygen–glucose deprivation (OGD) as previously described (50). Mice were euthanized by intraperitoneal Avertin injection and transcardially perfused with 20 mL cold sterile PBS. Spleens were harvested under sterile conditions and mechanically dissociated through a 70 μm cell strainer to generate a single-cell suspension. Red blood cells were lysed using Tris-ammonium chloride buffer (STEMCELL Technologies) for 10 minutes. Cells were then washed and resuspended in culture medium.

For OGD exposure, culture medium was replaced with serum-free, glucose-free Locke’s buffer (154 mM NaCl, 5.6 mM KCl, 2.3 mM CaCl₂, 1 mM MgCl₂, 3.6 mM NaHCO₃, 5 mM HEPES, and 5 mg/mL gentamicin; pH 7.2). Cells were placed in a hypoxia chamber maintained at 95% N₂ and 5% CO₂ for 1 hour. Control cells were maintained under normoxic conditions in glucose-containing medium at 95% air and 5% CO₂. Following OGD, splenocytes were treated with the pan-caspase inhibitor Q-VD-OPh (0.05 mM) or vehicle control for 1 hour prior to flow cytometry staining.

Cells were stained with Zombie Aqua viability dye (BioLegend) and incubated with Fc receptor blocking reagent (BioLegend) prior to staining with fluorophore-conjugated antibodies: CD45-R718, CD126 (IL-6R)-PE, Ly6G-eFluor450, CD11b-APC, and F4/80-PE-Cy7. Data were acquired using a CytoFLEX S (Beckman Coulter) or FACSMelody (BD Biosciences) and analyzed with FlowJo software (BD Biosciences). Myeloid populations were identified by sequential gating of CD45⁺ leukocytes followed by CD11b⁺ cells. Neutrophils were defined as CD11b⁺Ly6G⁺ cells, and macrophages as CD11b⁺F4/80⁺ cells. A minimum of 500,000 events were collected per sample. Fluorescence-minus-one (FMO) controls, tissue-matched controls, and unstained controls were used to establish gating thresholds.

#### Statistical Analysis

Data are presented as mean ± SEM with all individual data points shown, unless otherwise specified. Two-group comparisons were performed using unpaired t-tests with Welch’s correction. Comparisons involving more than two groups were analyzed by two-way ANOVA with Sidak’s multiple comparisons test. These analyses were applied to behavioral outcomes, brain atrophy measurements, ELISA assays (IL-6R, IL-6, LBP), and lung bacterial colony-forming unit (CFU) counts.

Survival data were analyzed using Kaplan–Meier survival curves with comparisons by the Mantel–Cox log-rank test. Neurological deficit scores (NDS) are presented as median (interquartile range) and were analyzed using two-way ANOVA with Sidak’s multiple comparisons test. Statistical significance was defined as p < 0.05. All mouse data analyses were performed using GraphPad Prism 9.

For human biomarker analyses, demographic variables were compared between sexes using the Wilcoxon rank-sum test for continuous variables and chi-squared or Fisher’s exact tests for categorical variables. Biomarker concentrations for IL-6, soluble IL-6 receptor (sIL-6R), and ADAM17 were log-transformed prior to analysis due to skewed distributions. Univariable comparisons were performed using two-sample t-tests. Multivariable linear regression models were used to assess associations between sex and biomarker levels, adjusting for age and NIH Stroke Scale (NIHSS) score. Human data analyses were conducted using R (version 4.5.2; R Foundation for Statistical Computing, Vienna, Austria).

#### Patient Samples

Patients with acute ischemic stroke, with confirmed images, admitted to Memorial Hermann Hospital (Houston, TX, USA) were enrolled in this study. All participants or their surrogates provided written or verbal informed consent (IRB-HSC-MS-17-0452). Plasma samples were collected 24 hours after the last known normal. All the patients had a MCAO stroke of more than 25% of MCA territory, and no hemorrhagic transformation. Exclusion criteria for all participants are recent blood transfusions, chronic steroid or immunosuppressant use, pregnancy, and active dialysis treatment. Detailed patient characteristics are provided in Supplementary Table 1A.

**Supplementary Table 1.**
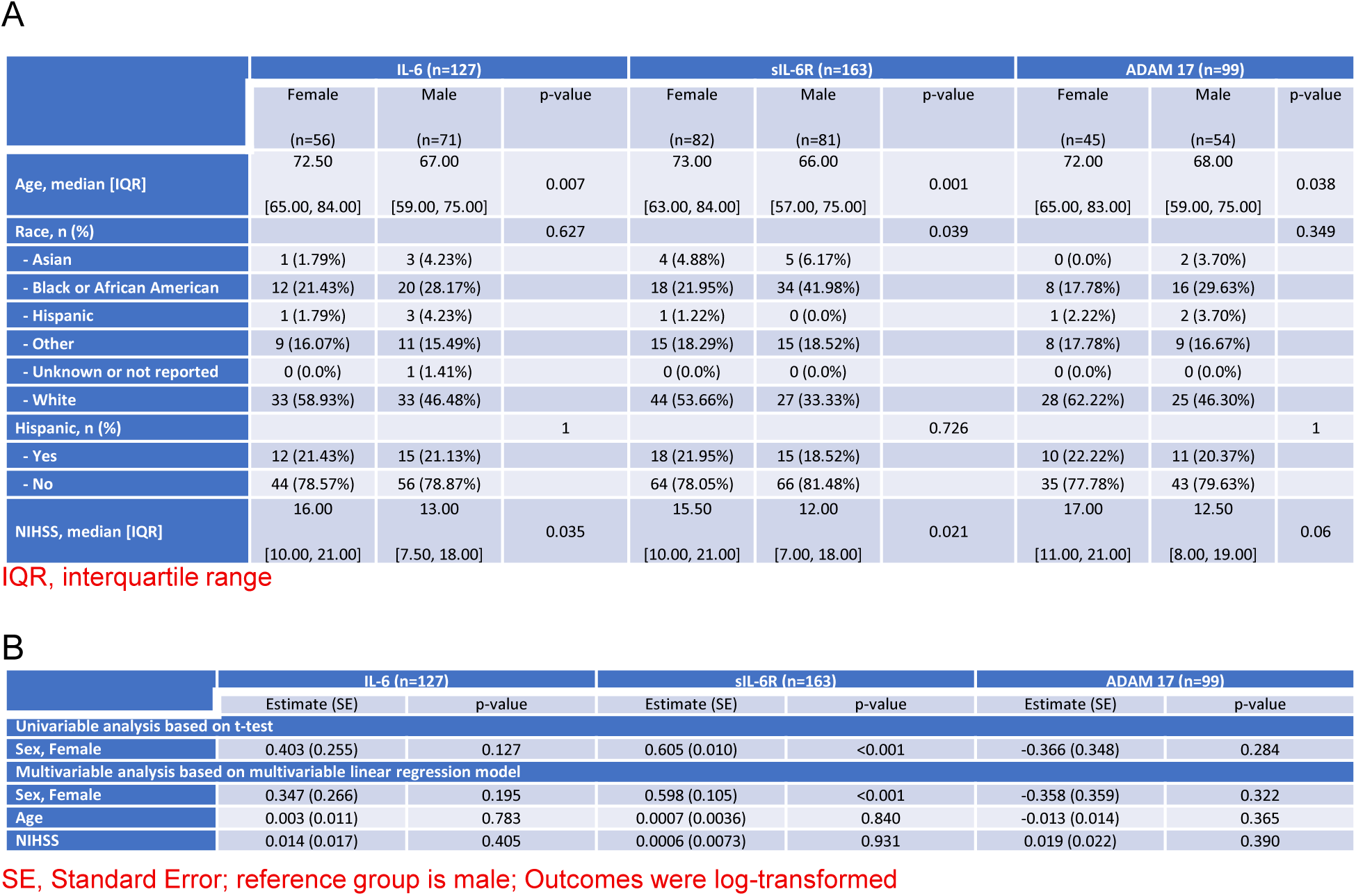
Patient demographics and analysis of circulating inflammatory markers. (A) Patient demographics stratified by sex.(B) Univariable and multivariable linear regression analyses of log-transformed IL-6, sIL-6R, and ADAM17 levels by sex.

**Supplementary Figure 1.**
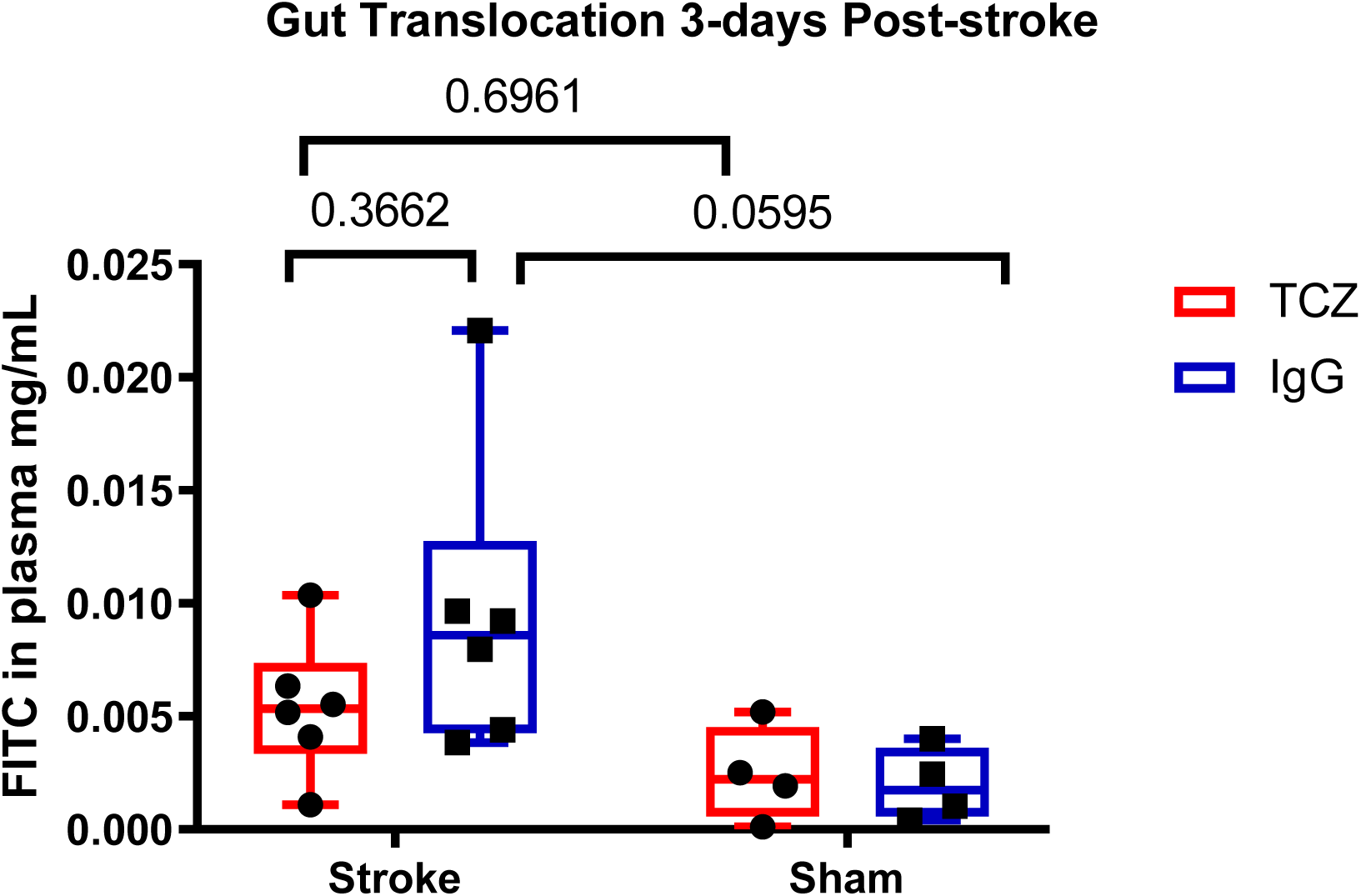
Gut translocation 3-days post-stroke. Intestinal permeability was assessed at 3-days post-stroke with fluorescein isothiocyanate-dextran (FITC-dextran), 4 kDa. Mice were fasted for 5-hours and then administered FITC-dextran via oral gavage (6mg/10g pre-stroke body weight) followed by euthanasia 1-hour later for measurement of plasma FITC-dextran concentration (n=4-6). Data presented as mean +/− SEM and analyzed using 2-way ANOVA with Tukey’s multiple comparisons test.

